# Competition and environmental gradients structure assemblages of Canidae across the planet

**DOI:** 10.1101/2021.01.09.426069

**Authors:** Lucas M. V. Porto, Rampal S. Etienne, Renan Maestri

## Abstract

The phylogenetic information of assemblages carries the signature of ecological and evolutionary processes that assembled these communities. Identifying the mechanisms that shape communities is not simple, as they can vary spatially. Here, we investigated how the phylogenetic structure of Canidae across the globe is affected by the environment and competition. We first identified phylogenetically clustered and overdispersed assemblages of canids over the planet taking into account regional pools of species in South and North America, Eurasia, and Africa. Then, we applied Structural Equation Models in these communities in order to identify the effect of temperature, vegetation cover, human impact, and body size dissimilarity on the spatial distribution of the phylogenetic information of canids. Only southern South America and the Middle East had strong phylogenetic clustered patterns, while the rest of the planet predominantly showed overdispersed assemblages. Body size dissimilarity and vegetation cover were the most important variables to explain the spatial patterns of phylogenetic structure in clustered and overdispersed communities, respectively. Canidae community composition across the world presents significant patterns of clustering and overdispersion, which vary following mainly the environmental gradient, suggesting habitat filtering as the main force acting on Canidae assemblages.

## 1. Introduction

The assembly of an ecological community is influenced by several biotic and abiotic factors (Webb 2000, Webb et al. 2002, Westoby 2006), including competition and habitat filtering. On the one hand, the competitive exclusion principle predicts that ecologically similar species cannot coexist if resources are limiting, and under phylogenetic trait conservatism, this implies that communities will be phylogenetically overdispersed (Darwin 1859, Elton 1946, Leibold 1998, Webb et al. 2002). On the other hand, phylogenetically closely related species likely share traits that allow them to tolerate a specific environment (Jarvinen 1982, Weiher et al. 1998, Webb et al. 2002, Di Marco and Santini 2015), which implies phylogenetic clustering of species in an ecological community. Because this also applies to traits that are linked to competitive ability, competition may also lead to phylogenetic clustering (Mayfield and Levine 2010, HilleRisLambers et al. 2012). If one can rule out this possibility, patterns of phylogenetic overdispersion or clustering are indicative of competition and habitat filtering, respectively, if the relevant traits are phylogenetically conserved. If traits are not conserved, overdispersion may be due to trait convergence of distant species, and clustering could be a result of historical processes that limited species’ dispersion from their ancestral ranges (Webb, 2000; Webb et al., 2002; Cavender-Bares et al., 2004; Kraft et al., 2007).

Webb (2000) and Webb et al. (2002) described how communities can be tested for phylogenetic overdispersion or clustering by comparing the value of a community metric to that of a null distribution of randomized communities. One such metric is the net relatedness index (NRI), where negative values of NRI are indicative of overdispersion, while positive values indicate clustering (Webb, 2000; Webb et al., 2002). They suggested that randomized communities can be obtained by permuting the presence-absence pattern on the phylogenetic tree. Pigot & Etienne (2015) argued that such a permutation approach does not yield a proper null distribution because it ignores speciation, colonization, and extinction dynamics. They developed a method that does take these processes into account and found that a null community (i.e. without habitat filtering or competition, but with speciation, colonization and extinction dynamics) would be overdispersed relative to the randomized community resulting from permutation. While absolute values for phylogenetic dispersions are therefore difficult to interpret, relative values are still informative: one can still compare dispersion patterns between communities and ask why some communities are more overdispersed or clustered than others. Many studies have used the phylogenetic approach described by Webb (2000) and Webb et al. (2002) to understand how the phylogenetic information of clades is structured through space (Cavender-Bares et al. 2004, Helmus et al. 2007, Kress et al. 2009, Kraft and Ackerly 2010, Goberna et al. 2014, Yang et al. 2014, Miazaki et al. 2015, Cadotte and Tucker 2017, Pérez-Valera et al. 2017, Zhang et al. 2018, Kusumoto et al. 2019). The majority of these studies have shown that overdispersion dominates at small scales, while clustering better explains the structure of communities at large scales. These findings demonstrate that the interpretation of the mechanisms acting in a community is scale-dependent (Webb et al. 2008). In this light, it is interesting to note that a large number of studies on phylogenetic structuring of communities are at small scales (see Cardillo, Gittleman, & Purvis, 2008).

Another difficulty in studies on the phylogenetic structure of communities relates to how abiotic and biotic factors are treated as independent forces acting on a community. Although habitat filtering and competition are contrasted in their effect on community structure, they occur together in natural communities (Ackerly 2003, Cadotte and Tucker 2017). Over the last decade, several studies have attempted to understand how much each mechanism influences communities (Kraft and Ackerly 2010, Goberna et al. 2014, Cadotte and Tucker 2017, Pérez-Valera et al. 2017, Zhang et al. 2018, Kusumoto et al. 2019). The results have been inconclusive. Cadotte & Tucker (2017) noted that “it is likely that most observational data reported as evidence for environmental filtering, in fact, reflect the combined effects of the environment and local competition.” To overcome these issues, one needs a detailed analysis of whether traits are conserved or not on a phylogenetic tree and of how patterns of phylogenetic dispersion vary with environmental variables.

Here, we assess patterns of phylogenetic dispersion in the family Canidae across the world. As canids are present in all continents, except in Antarctica (Wang and Tedford 2008, Wilson and Mittermeier 2009), and have a well-resolved phylogenetic tree (Porto et al., 2019), they are an ideal group to test how the phylogenetic structure of a whole clade was structured over the world by the influence of environmental filters and competition forces through time. The phylogeny of extant canids presents 36 species distributed in three distinct clades (Porto et al. 2019) with a variety of patterns: multiple species of a single genus, multiple genera of only one species, species with continental distributions, and also geographically restricted species. We explore, using Structural Equation Models (SEM), how phylogenetic dispersion patterns relates to environmental variables and to dissimilarity in body size (a measure of competition among canids).

## 2. Material and methods

### 2.1 Data compilation

Canids’ range maps were compiled from the IUCN Red List for all the 36 species (IUCN 2019). We processed these maps using ArcGIS (ESRI, 2019) to generate a presence-absence matrix of species within defined grids cells of 300 × 300 km (hereafter, assemblages). We used 3º grid cells because canids have very large home ranges. For each grid cell, we extracted average values for three environmental variables (mean temperature, vegetation cover, and human footprint) (Center for International Earth Science Information Network 2005, Fick and Hijmans 2017, USGS 2019). Temperature is the most significant environmental gradient at the global scale, and can strongly affect food-web structure and interaction strength of species (Bideault et al. 2019). Vegetation cover is suggested to have played a major role in the evolution of Canidae (Figueirido et al. 2015), and together with temperature, it constitutes two of the most commonly used variables in studies that evaluate species distributions (Porfirio et al. 2014). Human footprint was included in the models following Di Marco & Santini (2015), who found that human impacts explain the distribution of terrestrial mammals better than biological traits. Canid body sizes (head + body length) were obtained from the females of each species using the Handbook of the Mammals of the World (Wilson and Mittermeier 2009) and also from the Animal Diversity Web (ADW) (Myers et al. 2018) (Table S1). We used only the female body size to standardize the data due to sexual dimorphisms that a few species of canids present.

Other environmental variables were not taken into account for various reasons. Annual precipitation and primary productivity (Fick and Hijmans 2017) presented, within the assemblages addressed here (grid cells), high correlation with vegetation cover and temperature, respectively (Figure S1), which we already used in our models. The elevation gradient, which is commonly used in studies for different groups, was not considered here because only two species of Canidae have distributions that extend into mountains, so it is of little value in our study. Finally, including too many abiotic variables may bias the results toward environmental filtering.

We then calculated, for each grid cell, the body size dissimilarity among the canids within each assemblage using the decoupled trait approach proposed by De Bello et al. (2017) (*dcFdist*). This approach calculates functional differences between species accounting for the shared evolutionary history. Thus, we can measure how different canids within communities are, based on their body size, while accounting for their ancestry. Body size dissimilarity was used as a measure of competition. It is suggested by Finarelli (2007) that trends in body size were driven by interactions between species over the evolution of Canidae. One expects competition to be largest between species of similar body size, and if body size is evolutionarily conserved, competition causes phylogenetically related species to displace one another, so only phylogenetically distant species remain. Analyses were performed in R 4.0-2. (R Development Core Team 2020) using the raster package 3.3-13 (Hijmans and Etten 2012).

### 2.2 Phylogenetic data analyses

We used the phylogeny presented by Porto et al. (2019) that includes all 36 extant canid species and one recently extinct. The phylogeny was constructed through Bayesian inference based on 23 genes and 68 osteological characters. We removed the extinct species *Dusicyon australis* from this tree because there is no environmental data available for the region where the species lived before the year of its extinction (1876). To investigate how body size is distributed over the phylogeny, we calculated its phylogenetic signal using the *K*-statistic (Blomberg and Garland 2003) with the R package Phytools 0.7-47 (Revell 2012). Values of *K*□<□1 describe less phylogenetic signal than expected under a Brownian motion model of character evolution, while values of *K*□>□1 describe data with more phylogenetic signal than expected under Brownian motion.

To analyze phylogenetic dispersion patterns over the planet, we first calculated the standardized effect size of mean pairwise distances in communities (MPD) for each assemblage using the package Picante 1.8-2 (Kembel et al. 2010). As distributions of canids are not global, and some species are geographically restricted, historical processes (i.e. speciation and dispersal limitation) could influence our results. Thus, we set a null model with regional pools (instead of a global pool). We divided the world into 4 regional pools (South America = 10 species, North America = 9 species, Africa = 12 species, and Eurasia = 11 species); only few canids inhabit more than one continent (N. America + Eurasia: 2 species, Africa + Eurasia: 2 species, N. America + Eurasia + Africa: 1 species). Species richness spatial gradient and distributions are indicated in Figures S2 and S3, respectively.

Our null model randomized the community matrix (from each one of the four regions) 999 times by drawing canids from pool of species occurring in at least one community. Our null model shuffles taxa labels and re-calculates MPD, keeping richness fixed. The 999 null values constituted a null distribution to which the observed value was compared. We then calculated the net relatedness index (NRI) for the assemblages within each region using phylogenies containing only the species inhabiting those specific areas. We obtained the NRI from MPD as follows:

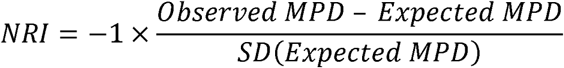

Larger (positive) NRI values imply a more phylogenetically clustered assemblage while smaller (negative) NRI values indicate more phylogenetic overdispersion. We chose the MPD to measure the relatedness of canids because it is probably the most popular measure for computing the phylogenetic distance between a given group of species which makes it easier for our results to be compared with the findings from other biological groups.

### 2.3 Structural equation models

To test the influence of biotic and abiotic variables on NRI patterns we used Structural Equation Models (SEM) (Mitchel 1992, Wang et al. 2013). SEM allow testing the contribution of different variables while accounting for potential correlations between them. We did not specify *a priori* whether any of these correlations had positive or negative effects. Our predictor variables (observed variables) were mean temperature, vegetation cover, human footprint, and body size dissimilarity. We assumed that all predictor variables affect NRI values and that there is a latent variable connected to our five observed ones to explain the covariation in the measurements. We tested four models (Figure 1), which differ as follows. Model 1 assumes that the human footprint has an impact on global temperature, which in turn affects vegetation cover. Model 2 is similar to model 1, but the human footprint affects vegetation cover, and the temperature is influenced by both variables. Neither of these two models have the influence of environmental variables on body size, but Models 3 and 4 do. In Model 3, body size dissimilarity is assumed to be influenced by vegetation and temperature, because higher values of these variables indicate more resources and hence may affect the level of competition which in turn can drive body size dissimilarity. Vegetation influences temperature in areas with high human density, but temperature also influences vegetation cover, for example, higher temperatures are associated with tropical forests. In Model 4, human footprint is assumed to have an effect on temperature and vegetation due to urbanization, and different from Model 3, there is no influence of vegetation on temperature.

**Figure 1.**
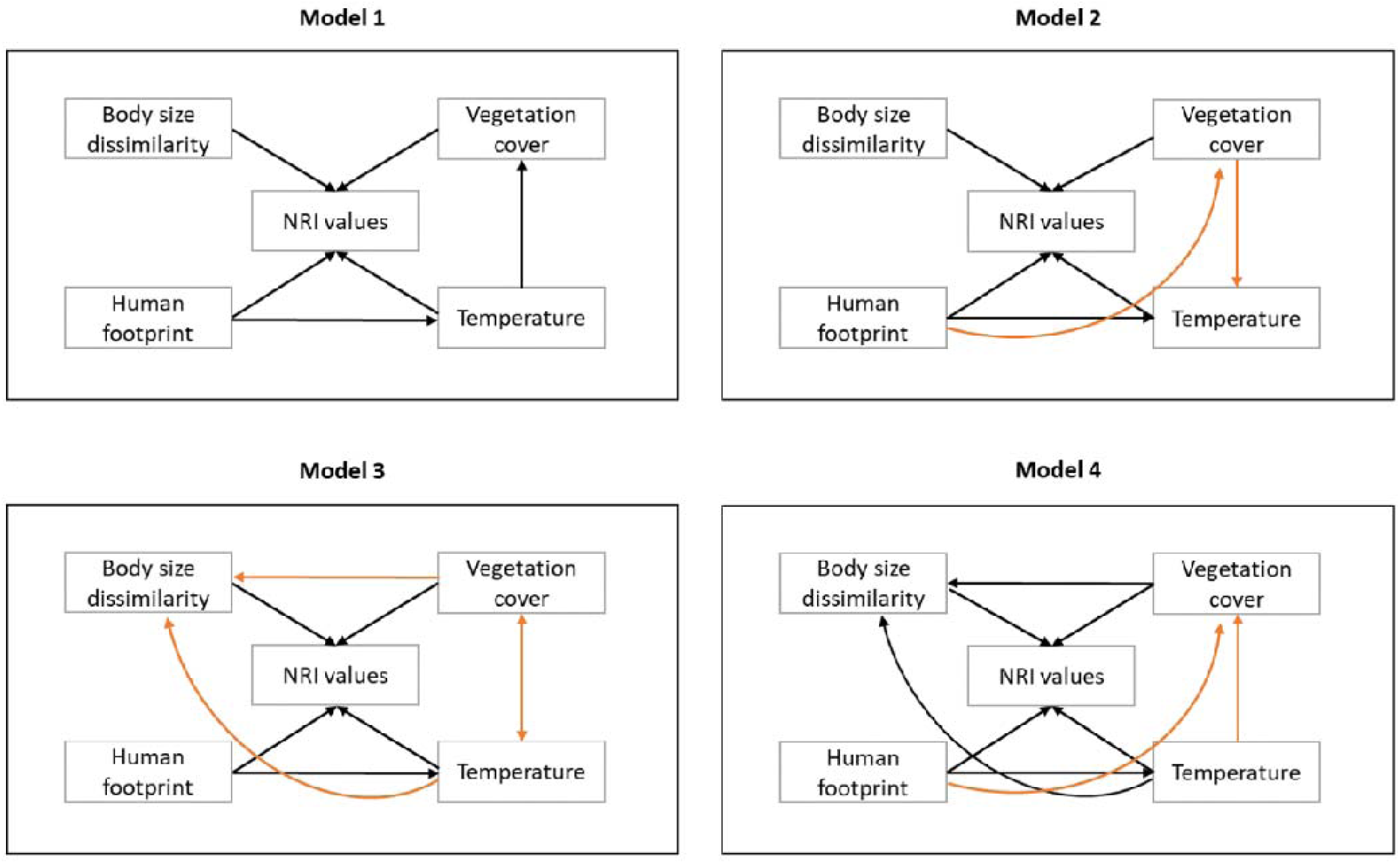
Schematic representations of the four Structural Equation Models tested in this study. The same models were used for communities with only negative, only positive, and all NRI values through separate analyses. Arrows indicate the direction of the effects of the variables. Orange arrows indicate different effects added to the model compared to the previous model. All models present a latent (hidden) variable connected to all five observed ones, but this is not represented here as it would unnecessarily pollute the images.

Because SEM analyses on all assemblages (phylogenetically overdispersed and clustered) simultaneously could mask some effect related to the predictor variables, we did not only analyze all data together, but we also conducted the SEM analyses separately for assemblages with negative NRI values and assemblages with positive NRI values. We conducted these analyses through maximum-likelihood estimation using the R package lavaan 0.6-7 (Rosseel 2012), and the models were compared using the Akaike information criterion (AIC) (Akaike 1973). Spatial autocorrelation was taken into account during the analyses using the package semTools 0.5-3 through the function “*spatialCorrect*” (Jorgensen et al. 2020). During the SEMs, the function takes a weighted matrix that reflects the intensity of the geographic relationship between observations and calculates Moran’s I for the residuals of all variables. If there is spatial autocorrelation in our dataset, the function spatially corrects it via Moran’s I. Another way to identify the mechanisms acting in communities would be applying the null model of assembly proposed by Pigot & Etienne (2015), DAMOCLES. This model considers the historical effects of speciation, colonization, and local extinction acting over time to determine the present composition of the community. However, DAMOCLES needs more species than we usually have in our communities to reliably estimate parameters.

## 3. Results

Phylogenetic signal in body size was high across the phylogeny of canids (*K* = 1.31, *p* < 0.01). The NRI values showed a dominance of overdispersed assemblages around the planet. Only southern South America and the Middle East had strong phylogenetic clustered patterns (Figure 2A). Asia presented a gradient of phylogenetic structure, as its southern and northern parts had clustered and overdispersed patterns, respectively, while its central region had no defined phylogenetic structure. Mean temperature, vegetation cover, human footprint, and body size dissimilarity also varied considerably across the world (Figures 2B, 2C, 2D, 2E).

**Figure 2.**
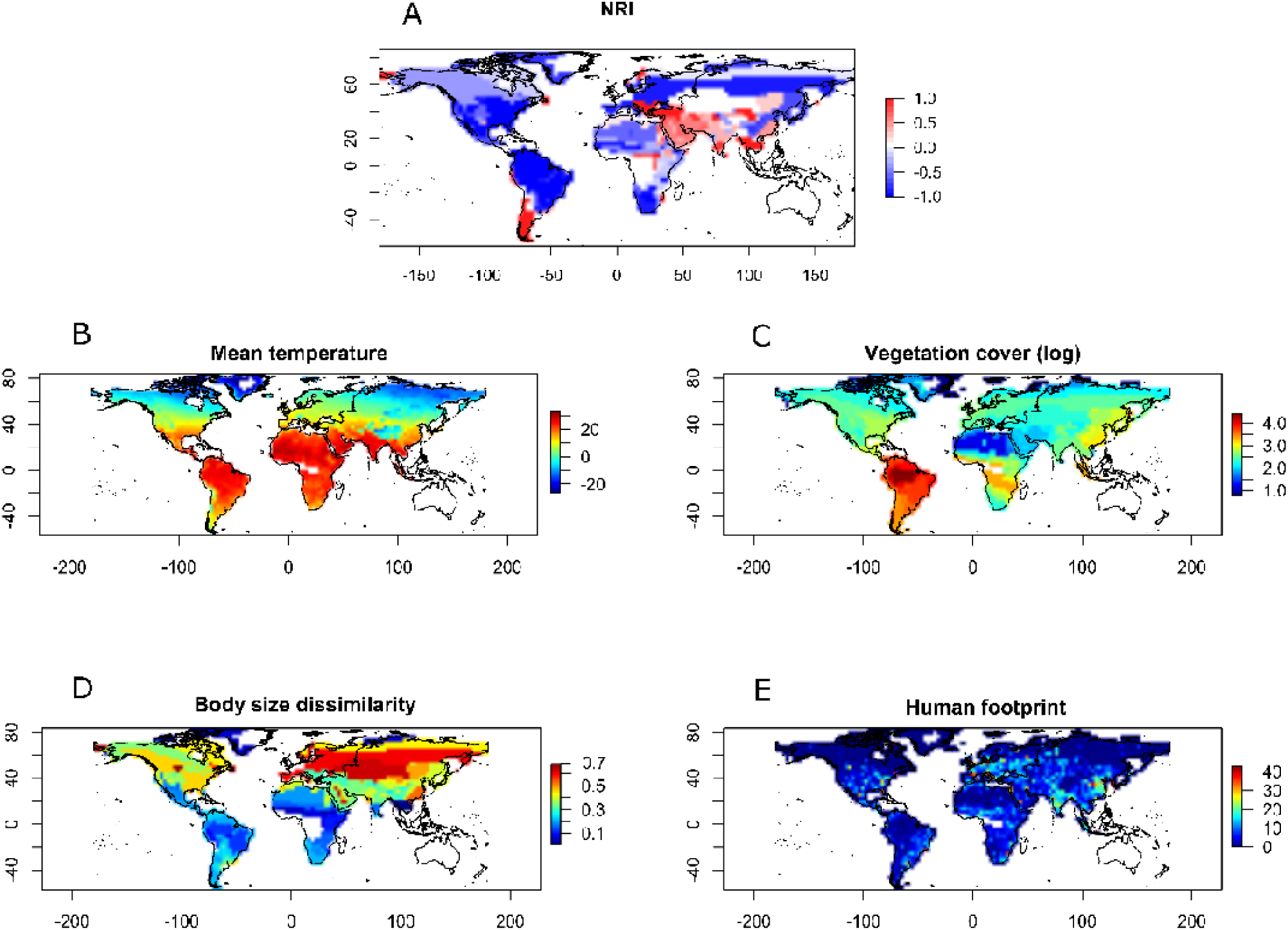
NRI and variables across the world. (A) NRI values for each assemblage showing that phylogenetic dispersion of canids varies spatially. Assemblages colored in blue present negative NRI values indicating that species within these areas are phylogenetically more dissimilar. Assemblages colored in red represent positive NRI values, where species are phylogenetically more similar. (B) Values of the global average temperature used in this study measured in Celsius (°C). (C) Global pattern of log-transformed vegetation cover percentage. (D) Body size dissimilarity (cm) among canids for each pixel. (E) Values of the human footprint (people per square kilometer).

Structural Equation Model 4 was selected as the best model for the three NRI analyses (negative NRI, positive NRI, and all NRI together), two AIC units better than the second-best model, model 3, in all cases (Table 1). The weights of model 4 were 0.831 for analyses with negative NRI, 0.863 for positive NRI and 0.744 for all NRI together. Model 3 presented weights of 0.168, 0.136 and 0.255 for negative, positive and all NRI together analyses, respectively. Models 1 and 2, with no influence of environmental variables on body size dissimilarity, performed poorly (weights less than 0.001). In Model 4, all predictor variables influenced NRI values, but also human footprint influenced vegetation and temperature, which in turn influenced vegetation cover and body size dissimilarity.

**Table 1.**
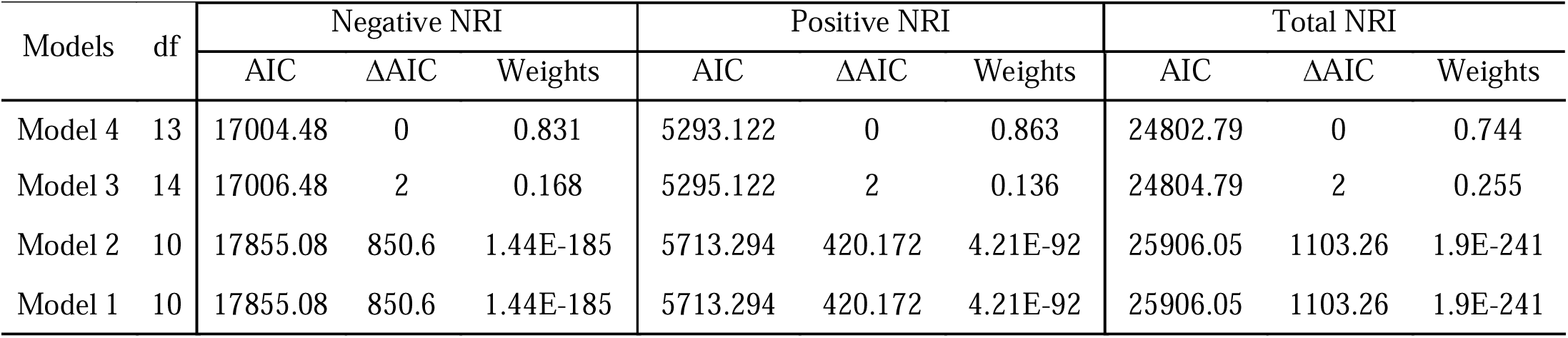
Akaike information criterion ranking of the four models of Structural Equation Models tested for negative, positive, and total NRI values.

| Models | df | Negative NRI |  |  | Positive NRI |  |  | Total NRI |  |  |
| --- | --- | --- | --- | --- | --- | --- | --- | --- | --- | --- |
|  |  | AIC | ΔAIC | Weights | AIC | ΔAIC | Weights | AIC | ΔAIC | Weights |
| Model 4 | 13 | 17004.48 | 0 | 0.831 | 5293.122 | 0 | 0.863 | 24802.79 | 0 | 0.744 |
| Model 3 | 14 | 17006.48 | 2 | 0.168 | 5295.122 | 2 | 0.136 | 24804.79 | 2 | 0.255 |
| Model 2 | 10 | 17855.08 | 850.6 | 1.44E-185 | 5713.294 | 420.172 | 4.21E-92 | 25906.05 | 1103.26 | 1.9E-241 |
| Model 1 | 10 | 17855.08 | 850.6 | 1.44E-185 | 5713.294 | 420.172 | 4.21E-92 | 25906.05 | 1103.26 | 1.9E-241 |

For all NRI data (Figure 3A), model 4 showed that the environmental variables temperature (*p* < 0.01, effect = 0.18) and vegetation (*p* < 0.01, effect = −0.33) were the most important to explain the spatial distribution of phylogenetic structure of canids (Table 2). Model 4 also showed that body size dissimilarity had a non-significant effect on NRI (*p* = 0.164), while human footprint had very weak influence on the phylogenetic structure of canids (*p* < 0.01, effect = 0.07).

**Table 2.** Results of the best-fitting Structural Equation Model (model 4) for all NRI values indicating the effect size among biotic and abiotic variables.

| Regressions: |  | Estimate | Std Err | P | Effect |
| --- | --- | --- | --- | --- | --- |
| NRI ~ | Body size dissimilarity | 0.119 | 0.085 | 0.164 | 0.038 |
|  | Human footprint | 0.007 | 0.002 | 0.003 | 0.079 |
|  | Temperature | 0.008 | 0.001 | < 0.01 | 0.188 |
|  | Vegetation cover | -0.014 | 0.001 | < 0.01 | -0.333 |
| Temperature ~ | Human footprint | 0.663 | 0.051 | < 0.01 | 0.321 |
| Vegetation cover ~ | Human footprint | -0.191 | 0.052 | < 0.01 | -0.093 |
|  | Temperature | 0.404 | 0.025 | < 0.01 | 0.408 |
| Body size dissimilarity ~ | Temperature | -0.005 | 0 | < 0.01 | -0.345 |
|  | Vegetation cover | -0.002 | 0 | < 0.01 | -0.177 |

\* *P* of the model (<0.01)

**Figure 3.**
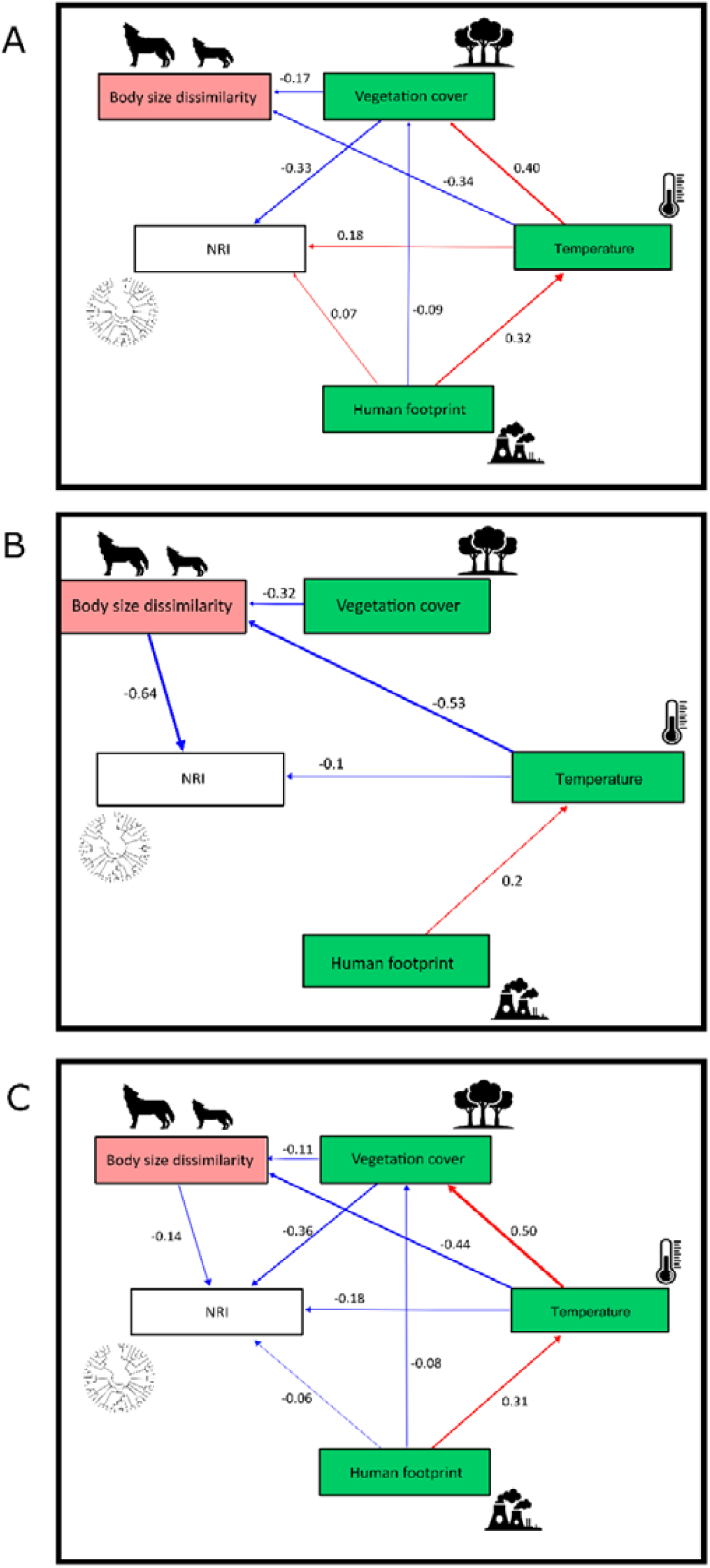
Best-fitting Structural Equation Model for the total NRI (A), clustered communities. i.e. with positive NRI (B), and overdispersed communities, i.e. with negative NRI (C). Positive and negative effects are indicated by red and blue arrows, respectively. Arrow thickness is scaled to illustrate the relative effect of each variable. Only significant effects (arrows) are shown (*p* < 0.01).

In the analysis on only the communities with positive NRI values, model 4 indicated that body size dissimilarity, our proxy for competition, was the most important variable to explain the phylogenetic composition within assemblages (*p* < 0.01, effect = −0.64) (Figure 3B, Table 3), indicating that the more dissimilar the communities in terms of body size, the less phylogenetically close the species within these communities were.

**Table 3.**
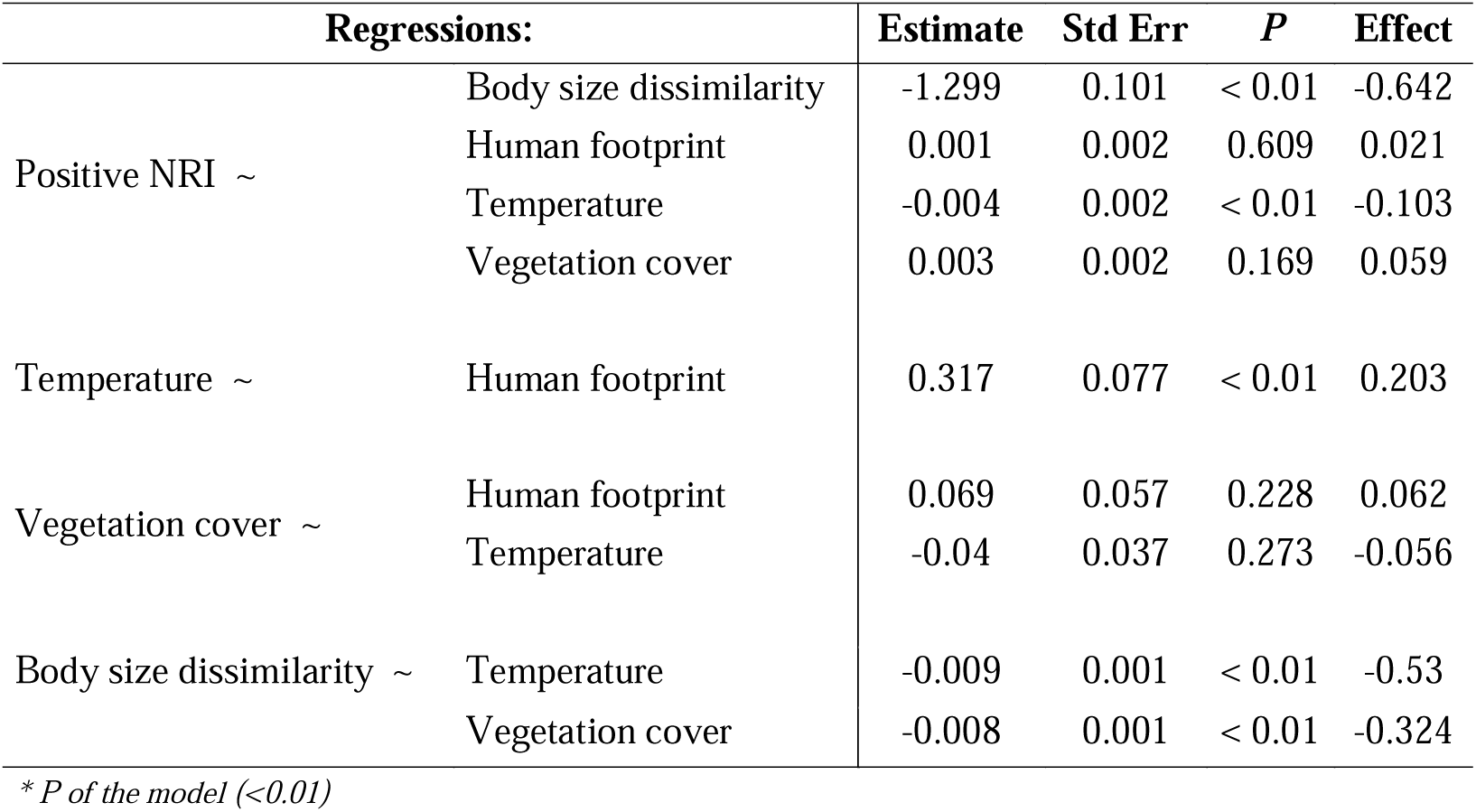
Results of the best-fitting Structural Equation Model (model 4) for positive NRI values indicating the effect size among biotic and abiotic variables.

In the analysis on only the communities with negative NRI values (Figure 3C), the environmental variables were the ones with the largest influence on phylogenetic dispersion. Based on model 4, vegetation cover was the most important variable (*p* < 0.01, effect = −0.36), suggesting that as vegetation cover increases, negative NRI decreases (becomes more negative), leading to species within these communities to become more phylogenetically distant (Table 4). Temperature (*p* < 0.01, effect = −0.18) and body size dissimilarity (*p* < 0.01, effect = −0.14) also had a considerable negative effect on NRI.

**Table 4.** Results of the best-fitting Structural Equation Model (model 4) for negative NRI values indicating the effect size among biotic and abiotic variables.

| Regressions: |  | Estimate | Std Err | P | Effect |
| --- | --- | --- | --- | --- | --- |
| Negative NRI ~ | Body size dissimilarity | -0.25 | 0.053 | < 0.01 | -0.144 |
|  | Human footprint | -0.003 | 0.001 | 0.036 | -0.06 |
|  | Temperature | -0.004 | 0.001 | < 0.01 | -0.188 |
|  | Vegetation cover | -0.007 | 0.001 | < 0.01 | -0.361 |
| Temperature ~ | Human footprint | 0.69 | 0.064 | < 0.01 | 0.313 |
| Vegetation cover ~ | Human footprint | -0.195 | 0.065 | < 0.01 | -0.084 |
|  | Temperature | 0.527 | 0.029 | < 0.01 | 0.503 |
| Body size dissimilarity ~ | Temperature | -0.005 | 0 | < 0.01 | -0.44 |
|  | Vegetation cover | -0.001 | 0 | < 0.01 | -0.11 |

\* P of the model (<0.01)

## 4. Discussion

The composition of canid communities across the world shows both negative and positive values of NRI and is influenced by distinct factors in different continents. However, it seems that, environmental filters played a greater role than traits related to biotic factors to explain the phylogenetic composition of assemblages of canids. This pattern was found in the analysis with all communities and in the analysis with negative NRI assemblages only (Figures 3A and 3C). Nevertheless, the biotic factor was very important in positive NRI assemblages (Figure 3B). This prominent effect could only be detected because we did the analyses separately, which supports the idea that it is necessary to separate the phylogenetic information (clustered and overdispersed patterns) to have a better understanding on how distinct variables structure assemblages through space.

The main factor structuring assemblages with negative NRI values was the vegetation cover, indicating that canids become less closely related as vegetation cover increases. However, the best model suggests that overdispersed communities are maintained due to the combined effect of vegetation cover, temperature, and body size dissimilarity (Figure 3C).

Interestingly, the relationship between body size dissimilarity and NRI had the same direction in positive and negative NRI communities. In clustered communities, species were less phylogenetically close as body size dissimilarity increased, while in overdispersed communities, phylogenetically distant species were associated with larger body size dissimilarity. Note that body size dissimilarity has been corrected for phylogenetic signal, so this body size variation was more than can be expected from shared evolutionary history. Hence, this implies that species are much more dissimilar in their body sizes at very negative NRI values (highly overdispersed communities) and more similar in their body sizes at very positive NRI values (highly clustered communities).

The relationship between body size dissimilarity and NRI might be explained by two distinct processes. In phylogenetically clustered communities our proxy for competition was the variable with the strongest influence on NRI values, but this pattern may actually be controlled by an environmental filter (which is the force we would expect to be dominant in communities with species that are phylogenetically more closely related than expected by chance), because temperature and vegetation cover had strong effects on body size dissimilarity, and therefore competition seems to be unlikely as the main force in these communities. Thus, body size must have played an important role in species persistence under the effect of some filter generated by both environmental variables. By contrast, in overdispersed communities competition may not have influenced these regions due to the low association between body size dissimilarity and NRI, but if it acted on these assemblages it may be explained by competition after secondary contact, because in overdispersed communities there are no longer such strong effects of environmental variables on body size. These explanations assume that clustered communities undergo mostly sympatric speciation whereas overdispersed communities assembled through immigration after allopatric speciation. This seems to be in line with the results from Porto et al. (2021), on the origin and dispersal of the Canidae lineages. They show that highly clustered communities, such as the Middle East and South America, present several lineages that originated within these regions. Furthermore, highly overdispersed regions, such as North America, Europe and South Africa, are inhabited by lineages of distinct clades, where many have their origins outside these regions.

The overdispersed pattern presented by assemblages of canids living in North America, Europe, and the northern part of Asia was probably due to a large number of distinct species of the genera *Vulpes, Canis*, and *Urocyon* that coexist within these areas and are very phylogenetically distant, being separated around 12 million years ago (Ma) (Porto et al. 2019). Species from these three genera have very distinct diet, habitat type, and social behavior (Wang and Tedford 2008, Wilson and Mittermeier 2009).

For phylogenetically clustered assemblages, human footprint did not present any significant influence on NRI values (Figure 3B), and only a very small effect in both other sets of analyses (Figure 3A and 3C). This is surprising because Di Marco & Santini (2015) demonstrated that human impacts influence the geographical distribution of terrestrial mammals better than the biological traits of species. Anthropogenic impacts seem to be important to understand the spatial distribution of the phylogenetic information of Canidae, considering the reports of endemic canids in the Americas constantly losing habitat and being killed by humans (Hoffmann et al. 2011), but our analyses do not support this.

We emphasize the better performance that models with environmental variables influencing body size dissimilarity had compared to the ones that did not. This suggests environmental control of body size dissimilarity. For overdispersed assemblages, as temperature decreases, dissimilarity increases, suggesting greater competition in colder regions of the planet. In clustered communities, temperature strongly affects body size dissimilarity too, showing that as temperature increases, species become similar in their sizes (less dissimilarity), suggesting that warm regions within clustered assemblages, such as deserts and tropical forests, have imposed a much greater environmental filter on canids than cold regions. Our finding is contrary to what was proposed by Dobzhansky (1950), who argued that biotic factors are more limiting in the tropics, while abiotic conditions are more important at higher latitudes. However, our finding may not be so surprising given that the first ancestors of the living canids were probably packing hunters with medium body sizes, around 70 cm (Wang and Tedford 2008, Porto et al. 2019). Deserts and tropical forests may act as a strong filter to canids with such a lifestyle because food is scarce in deserts and pack hunting is difficult inside dense forests. An alternative explanation may be differential diversification rates. Pigot & Etienne (2015) showed that allopatric speciation creates overdispersed patterns, suggesting that for canids speciation might be higher at higher latitudes. Although SEMs in clustered and overdispersed communities indicated a negative correlation between NRI and temperature, this effect is positive when all-NRI values are analyzed together. We are aware of this apparent contradiction, but it makes clear that splitting the data in positive and negative NRI reveals more direct relationships whereas relationships are more indirect in the all-NRI scenario.

Even though canids are not very dispersal limited, because they can travel long distances (Wang and Tedford 2008, Wilson and Mittermeier 2009), they may still tend to be found in higher numbers near their center of diversification than far from it. This can also help to understand the phylogenetic structure of the Middle East and the southern portion of South America, which, based on the fossil records and biogeographical models, had major diversification events of foxes and South American canids, respectively (Wang and Tedford 2008, Porto et al. 2021). Ecological speciation within these regions might have generated species’ phylogenetic clustering. In these areas the number of endemic species is high, and speciation tends to generate similar trait values (Gillespie 2004).

The majority of studies of the phylogenetic structure within communities concerns plants or focus on small geographic scales. Nevertheless, some studies with vertebrates at large spatial scales have demonstrated contrasting patterns of phylogenetic composition. Cooper, Rodríguez, & Purvis, (2008) found a tendency of overdispersion in assemblages of New world monkeys, Australasian possums, and North American ground squirrels. Yan et al. (2016), however, found phylogenetic clustering for Mammalia, Aves, Reptilia, and Amphibia from China. And Cardillo (2011) demonstrated an unstructured phylogenetic pattern on African carnivore’s assemblages. Here we presented a case where both patterns are important to understand community composition across the planet, depending on the region studied.

Larger geographical scales are expected to generate more phylogenetic clustering than overdispersion because the rate of speciation increases with more space available (Losos and Schluter 2000) due to higher habitat heterogeneity (Kneitel and Chase 2004), and thus sister species are more likely to co-occur at larger scales. Even though we found some areas with phylogenetic clustering (Figure 2A), there was a dominance of overdispersed communities, indicating that the scale we used (300 km × 300 km) may still be too small for clustering to kick in. Therefore, we anticipate that using an even larger scale, e.g. the biome scale, will reveal clustering. Studying community structure at that scale, however, is no longer very informative, exactly because of the large expected biogeographic contribution to phylogenetic dispersion.

Sandel (2018) suggested that, if there is substantial variation in species richness along the geographical gradient, the effects that biotic and abiotic factors have on community assembly should not be assessed by only comparing index values, because NRI may depend on species richness. However, this is not the case here, as the species richness spatial gradient for Canidae does not present considerable variation over the planet (Figure S2).

The mechanisms that influence the assembly of a community can act in complex ways (Ricklefs 1987, 2015). In this study we used both a phylogenetic approach and an approach based on environmental variables and traits. We demonstrated that canid community composition across the world presents substantial patterns of clustering and overdispersion that follow mainly the environmental gradient, suggesting habitat filtering as the main force acting on Canidae assemblages.

## Acknowledgements

We thank Vanderlei Debastiani for the help and suggestions during the analysis. This study was financed in part by the Coordenação de Aperfeiçoamento de Pessoal de Nível Superior - Brazil (CAPES) and by the University of Groningen. RSE thanks the Dutch Research Council (NWO) for funding through a VICI grant.

## Declaration section

### Ethics approval and consent to participate

Not applicable.

### Consent for publication

Not applicable.

### Availability of data and material

All data generated or analyzed during this study are included in this published article.

## Authors’ contributions

L.M.V.P. conceived and designed the study and analyses. L.M.V.P. performed the analyses. L.M.V.P. wrote the first draft of the manuscript. R.S.E. and R.M. commented on the methods and contributed to substantial revisions on the draft.

## Supplementary material

**Figure S1.**
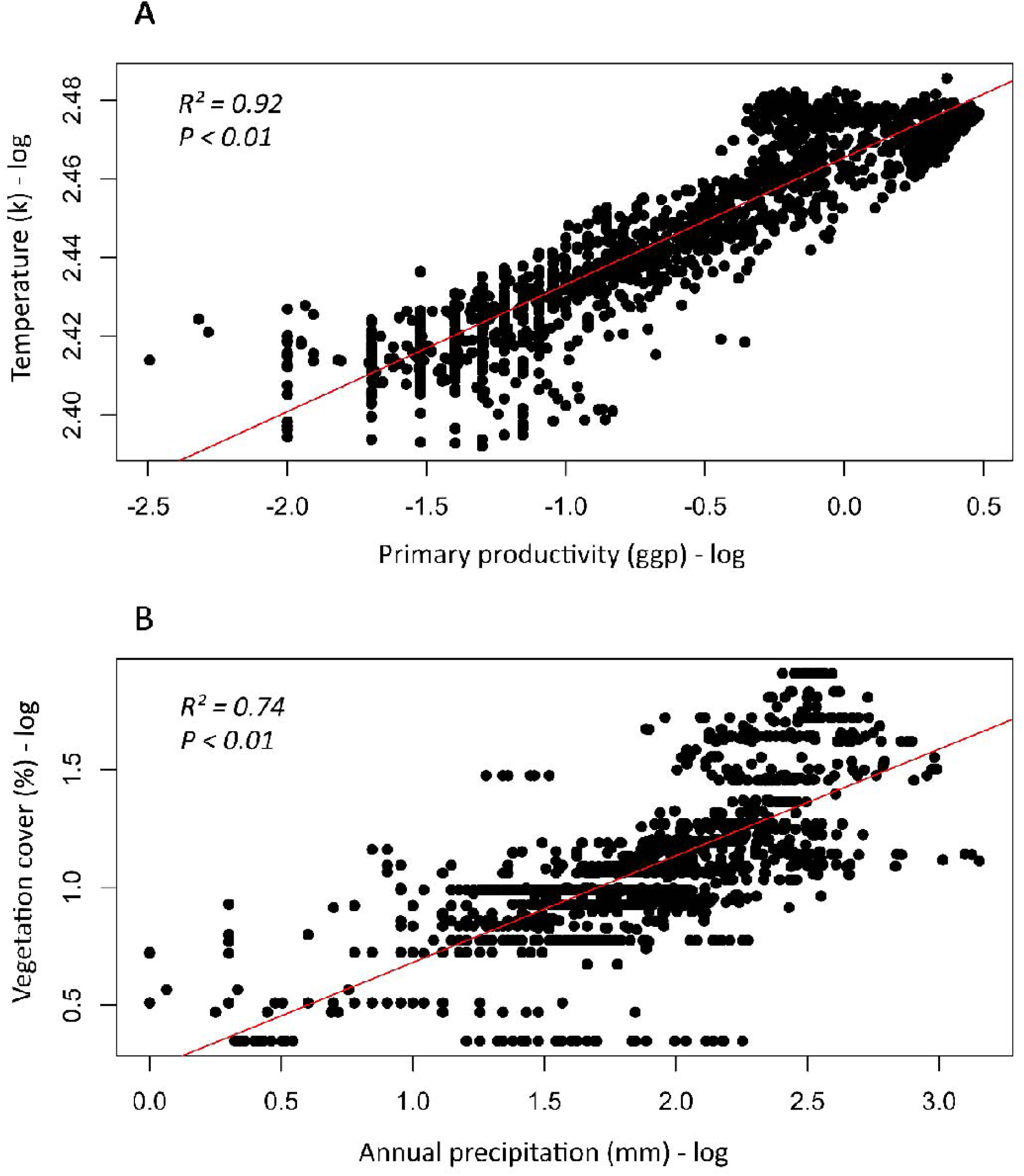
(A) Correlation between annual temperature and primary productivity (R^2^ = 0.92, *p* < 0.01). (B) Correlation between vegetation cover and annual precipitation (R^2^ = 0.74, *p* < 0.01). Only values extracted from the assemblages used here are represented in the correlations.

**Figure S2.**
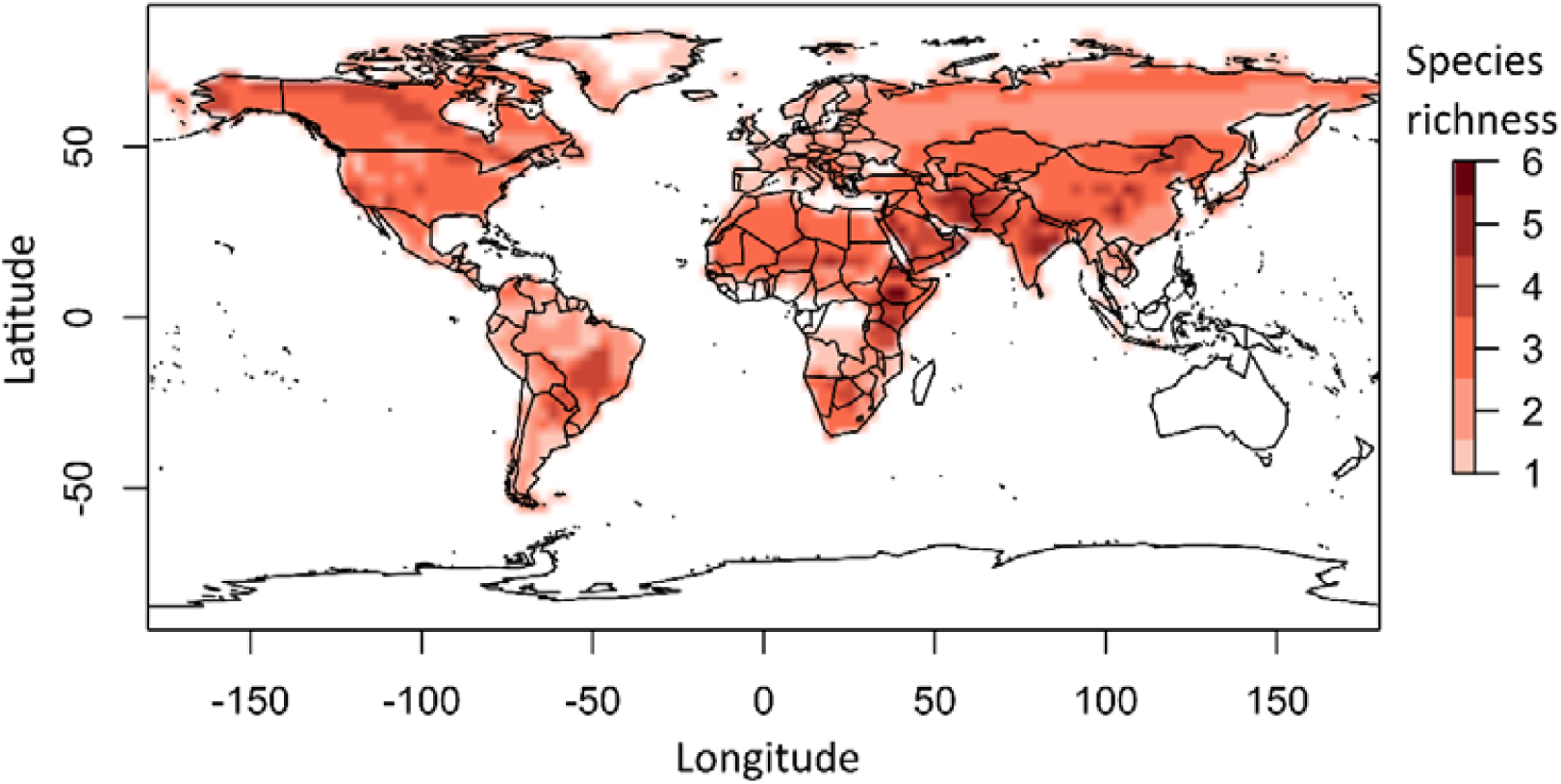
Species richness spatial gradient for all 36 Canidae species in this study.

**Figure S3.**
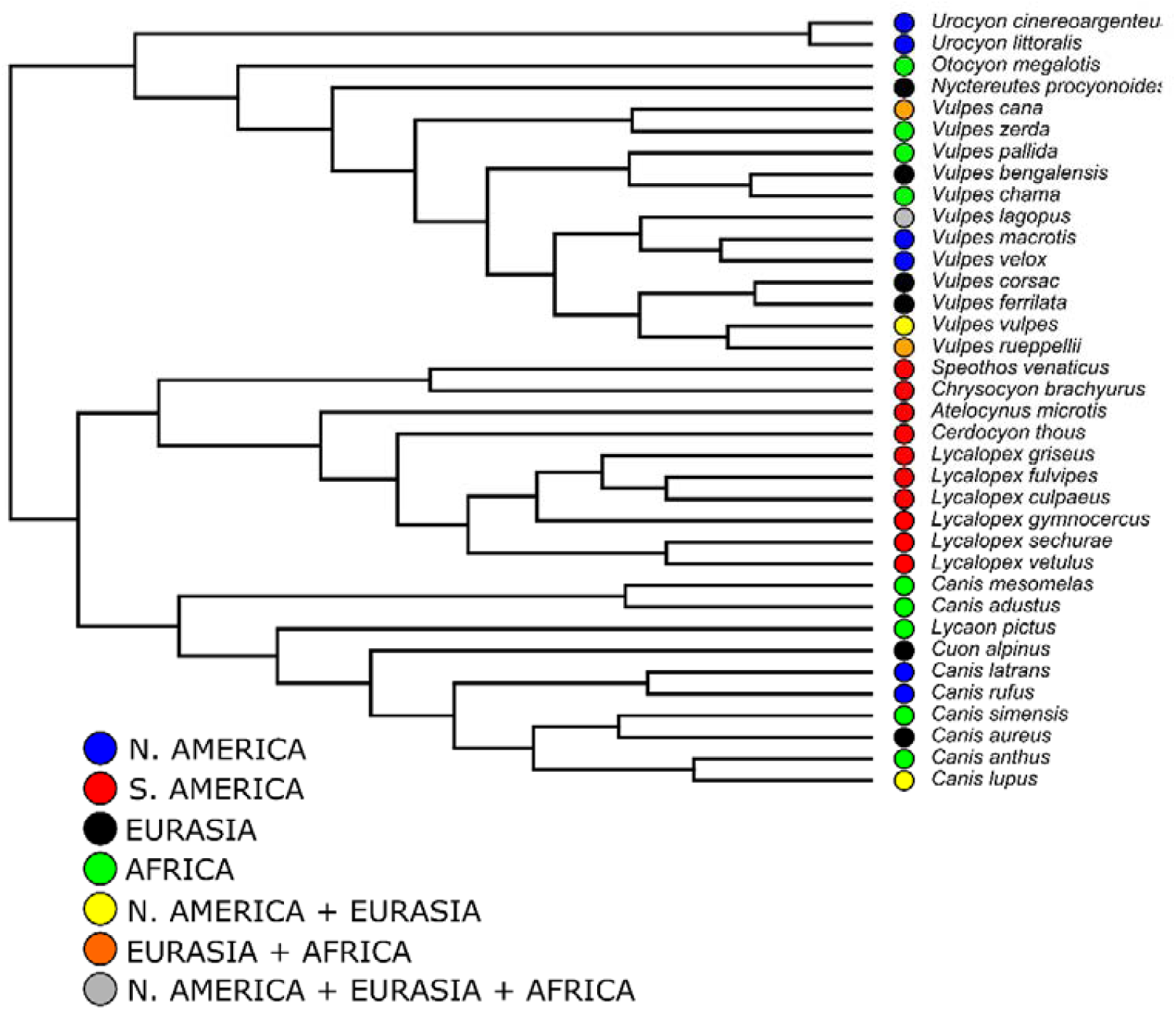
Species distributions indicated along the tips of the phylogeny from Porto et al. (2019) that was used in our study.

**Table S1.**
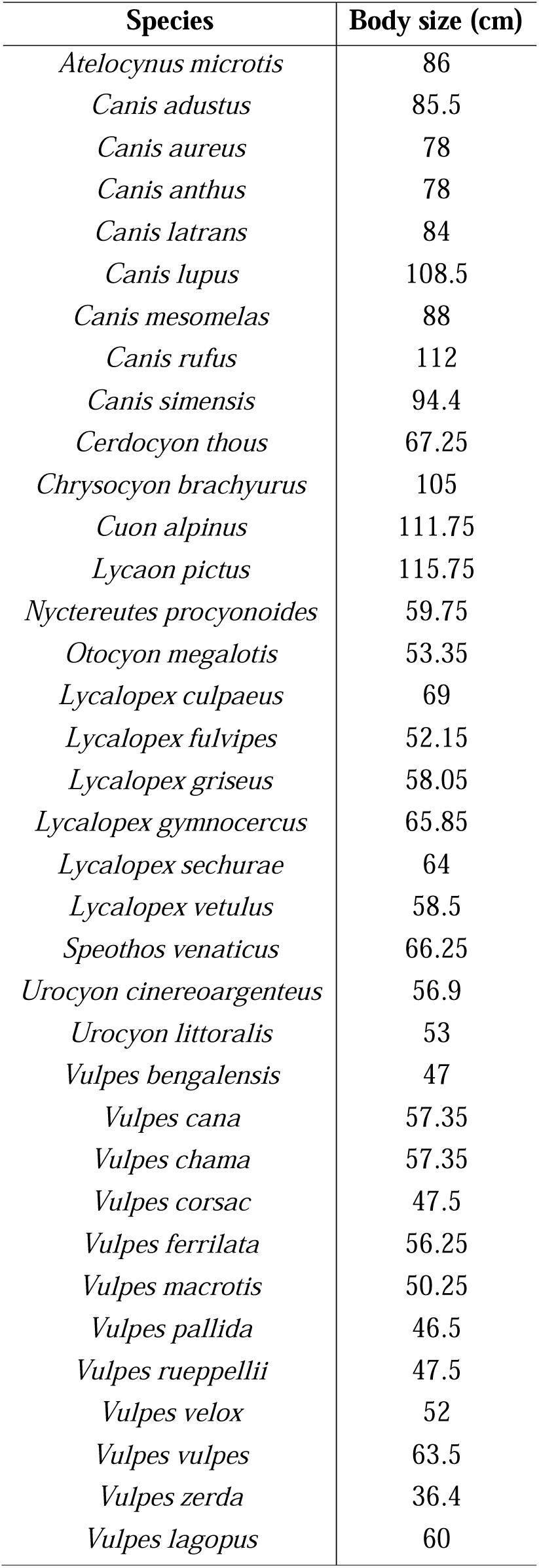
Body size of the 36 species of canids used in this study. The body size measurements used here are actually the head + body length measurements obtained from the females of each species.

| Species | Body size (cm) |
| --- | --- |
| *Atelocynus microtis* | 86 |
| *Canis adustus* | 85.5 |
| *Canis aureus* | 78 |
| *Canis anthus* | 78 |
| *Canis latrans* | 84 |
| *Canis lupus* | 108.5 |
| *Canis mesomelas* | 88 |
| *Canis rufus* | 112 |
| *Canis simensis* | 94.4 |
| *Cerdocyon thous* | 67.25 |
| *Chrysocyon brachyurus* | 105 |
| *Cuon alpinus* | 111.75 |
| *Lycaon pictus* | 115.75 |
| *Nyctereutes procyonoides* | 59.75 |
| *Otocyon megalotis* | 53.35 |
| *Lycalopex culpaeus* | 69 |
| *Lycalopex fulvipes* | 52.15 |
| *Lycalopex griseus* | 58.05 |
| *Lycalopex gymnocercus* | 65.85 |
| *Lycalopex sechurae* | 64 |
| *Lycalopex vetulus* | 58.5 |
| *Speothos venaticus* | 66.25 |
| *Urocyon cinereoargenteus* | 56.9 |
| *Urocyon littoralis* | 53 |
| *Vulpes bengalensis* | 47 |
| *Vulpes cana* | 57.35 |
| *Vulpes chama* | 57.35 |
| *Vulpes corsac* | 47.5 |
| *Vulpes ferrilata* | 56.25 |
| *Vulpes macrotis* | 50.25 |
| *Vulpes pallida* | 46.5 |
| *Vulpes rueppellii* | 47.5 |
| *Vulpes velox* | 52 |
| *Vulpes vulpes* | 63.5 |
| *Vulpes zerda* | 36.4 |
| *Vulpes lagopus* | 60 |

